# A Sequence-based Antibody Paratope Prediction Model Through Combing Local-Global Information and Partner Features

**DOI:** 10.1101/2021.07.17.452531

**Authors:** Shuai Lu, Yuguang Li, Xiaofei Nan, Shoutao Zhang

## Abstract

Antibodies are proteins which play a vital role in the immune system by recognizing and neutralizing antigens. The region on the antibody binding to an antigen, known as paratope, mediates antibody-antigen interaction with high affinity and specificity. And the accurate prediction of those regions from antibody sequence contributes to the design of therapeutic antibodies and remains challenging. However, the experimental methods are time-consuming and expensive. In this article, we propose a sequence-based method for antibody paratope prediction by combing local and global information of antibody sequence and partner features from partner antigen sequence. Convolution Neural Networks(CNNs) and a sliding window approach on antibody sequence are used to extract local information. Attention-based Bidirectional Long Short-Term Memory (Att-BLSTM) on antibody sequence are used to extract global information. Also, the partner antigen is vital for paratope prediction, and we employ Att-BLSTM on the partner antigen sequence as well. The outputs of CNNs and Att-BLSTM networks are combined to predict antibody paratope by fully-connected networks. The experiments show that our proposed method achieves superior performance over the state-of-the-art sequenced-based antibody paratope prediction methods on benchmark datasets.

## 1 Introduction

A NTIBODIES play an import role in human immune system by binding a wide range of macromolecules with high affinity and specificity[1]. This ability of binding is mediated by the interaction of amino acid residues at the paratope region of antibody[2]. Accurate prediction of antibody paratope is helpful for the study of antibody-antigen interaction mechanism and the development of antibody design[3]–[5]. Along with the establishment of antibody expression and purification process, the number of approved therapeutic antibodies is growing steadily[6]. As a result, correct judgement of which amino acid residues belonging to paratope is essential[7].

The antibody paratope region is usually determined by observing the residues that are spatially close to its partner antigen from its structure obtained by X-ray[8], NRM[9] and Cryo-EM[10]. However, these experimental methods are time-consuming and expensive[11]. Therefore, a lot of computational models are developed to overcome these problems. According to the input features, the computational methods for antibody paratope prediction can be classified into two classes: sequence-based and structure-based. As the name implies, the sequence-based models only use the antibody sequence features and the structure-based models utilize the antibody structure features as well. Some of the sequence-based methods take the whole antibody primary sequence as input and output the binding probability of each amino acid residue[12]–[14]. Other methods only utilize the residues in complementarity determining regions (CDRs) and predict their binding probability[15], [16]. However, about 20% of the residues that participating in binding fall outside the CDRs[17]. Most structure-based methods consider antibody structure as a graph and employ graph convolution operation for aggregating the information of spatial neighboring residues[18]–[20]. Apart from those methods, 3D Zernike descriptors are used for representing a set of amino acid residues adjacent to each other, and shallow machine learning models are used[21]. Although structure-based models can provide more accurate description of paratope, sequence information is always available earlier than structure[15].

Global information of the whole sequence has been proved useful in many biology biological analysis tasks such as protein-protein interaction sites prediction[22] and protein phosphorylation sites prediction[23]. Methods from those works extract global features form the whole protein sequence by TextCNNs[24] or SENet blocks[25] and Bi-LSTM blocks[26]. In this study, we utilize Attention-based Bidirectional Long Short-Term Memory(Att-BLSTM) to extract global features from antibody sequence and partner features form its partner antigen sequence, as Att-BLSTM shows superior performance in several machine learning tasks[27], [28].

In this work, we propose a sequence-based method for antibody paratope prediction utilizing both local-global information and partner features by combing Convolutional Neural Networks (CNNs) and Att-BLSTM networks. For local features, we use CNNs with a fixed sliding window size to consider the neighbor information around a target amino acid residue on antibody sequence. For global features and partner features, we use Att-BLSTM networks on antibody sequence and its partner antigen sequence, respectively. After that, all features are combined to fed into fully-connected networks to predict the probability of each antibody residue belonging to paratope. We also compare results with other competing sequence-based paratope predictors, and our method achieves the best performances.

## 2 Materials and methods

### 2.1 Datasets

The datasets used in this study are taken from PECAN[20]. All complexes are filtered to make sure that no antibody sequence share identity more than 95%. The complexes in training and validation sets are collected from the training sets of other paratope prediction works[12], [15], [29] and AbDb[30]. The training set contains 205 complexes and the validation set consists of 103 complexes to tune the hyper parameters in our model. And, 152 complexes are used for testing and evaluating the performance of our model.

Similar as other works, a residue on antibody is defined to belong to paratope if any of its heavy atoms is less than 4.5 Å away from any heavy atom on antigen[15], [20], [29]. It should be noted that the structure of antibody and antigen complex is used only extracted positive and navigate labels of antibody residues. The summary of datasets is shown in Table 1.

**TABLE 1.** Number of complexes and residues in the datasets.

| DataSet | Complexes | Positive residues | Negative residues |
| --- | --- | --- | --- |
| Training | 205 | 4449 (5.19%) | 81283 (94.81%) |
| Validation | 103 | 2237 (5.24%) | 60584 (94.76%) |
| Testing | 152 | 3314 (5.19%) | 40480 (94.81%) |

### 2.2 Input features

We utilize residue features consist of one-hot encoding, evolutionary information, seven additional parameters and predicted structural features of antibody or antigen sequence. All of them are described in details as follows:

#### 2.2.1 One-hot encoding of antibody sequence

Only 20 natural types of amino acid residues are considered in our study. Each type is encoded to a 20D vector where each element is either 1 or 0 and 1 indicates the existence of a corresponding amino acid residue.

#### 2.2.2 Evolutionary information

A lot of studies use the evolutionary information of antibody or protein sequence for biological analysis tasks, such as B-cell epitope prediction[31], [32], protein-protein interaction sites prediction[22], [33]–[36], protein-DNA binding residues prediction[37], protein folding recognition[38] and protein contact map prediction[39]. In this study, we run PSI-BLAST[40] against the nonredundant (nr)[41] database for every antibody and antigen sequence in our datasets with three iterations and an E-value threshold of 0.001. After that, we get the position-specific scoring matrix (PSSM) and the position-specific frequency matrix (PSFM). Each amino acid is encoded as a 20D vector representing the probabilities and frequencies of 20 natural amino acid residues occurring at each position in PSSM and PSFM, respectively. For each protein sequence with L residues, there are L rows and 20 columns in PSSM or PSFM. Besides PSSM and PSFM, two parameters at each residue position are obtained as well. One is related with column entropy in multiple sequence alignment, and another is related with column gaps in multiple sequence alignment.

#### 2.2.3 Seven additional parameters

Those parameters represent physical, chemical and structural properties of each type of amino acid residue by training artificial neural networks for protein secondary structure prediction[42].

#### 2.2.4 Predicted structural features

In this study, we predict antibody or antigen local structural features from sequence by NetSurfP-2.0 which is a novel deep learning model trained on several independent datasets[43]. Solvent accessibility, secondary structure, and backbone dihedral angles for each residue of the input sequences are returned from NetSurfP-2.0. Among those features prediction tasks, NetSurfP-2.0 achieves the state-of-the-art performance.

We calculate the absolute and relative solvent accessibility surface accessibility(ASA and RSA, respectively), 8-class secondary structure classification (SS8), and the backbone dihedral angles(*ϕ* and *ψ*) for each residue position of input antibody or antigen sequence. ASA and RSA represent the solvent accessibility of an amino acid residue. The predicted secondary structure describes the local structural environment of a residue. And, *ϕ* and *ψ* figure the relative positions of adjacent residues. The 8-class secondary structures are: 3-helix (G), a-helix (H), p-helix (I), b-strand (E), b-bridge (B), b-turn (T), bend (S) and loop or irregular (L). And, we use the one-hot encoding of SS8.

Together, an 80D vector is used for representing a residue in our study.

### 2.3 Input representation

The antibody paratope prediction problem can be summarized as a binary classification task: judging whether a residue from a given antibody sequence binding with its partner antigen or not. As described in Section 2.2, each residue is encoded into an 80D vector. And each antibody or antigen sequence can be represented as a matrix *S*, including a list of residues:

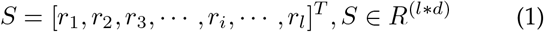

where *r*_*i*_ *∈ R*^*d*^ is the residue feature vector corresponding to the *i*-th residue in the antibody or antigen sequence, *l* is the sequence length, and *d* is the residue feature dimension(80 in this paper).

### 2.4 Model architecture

As Fig.1 shows, our proposed method is mainly composed of three parallel parts: CNNs extract local features from antibody sequence, Att-BLSTM networks extract global features from antibody sequence and another Att-BLSTM network extracts partner features from partner antigen sequence. The outputs of those parts are concatenated and fed to fully connected networks to predict the binding probability for each antibody residue.

**Fig. 1.**
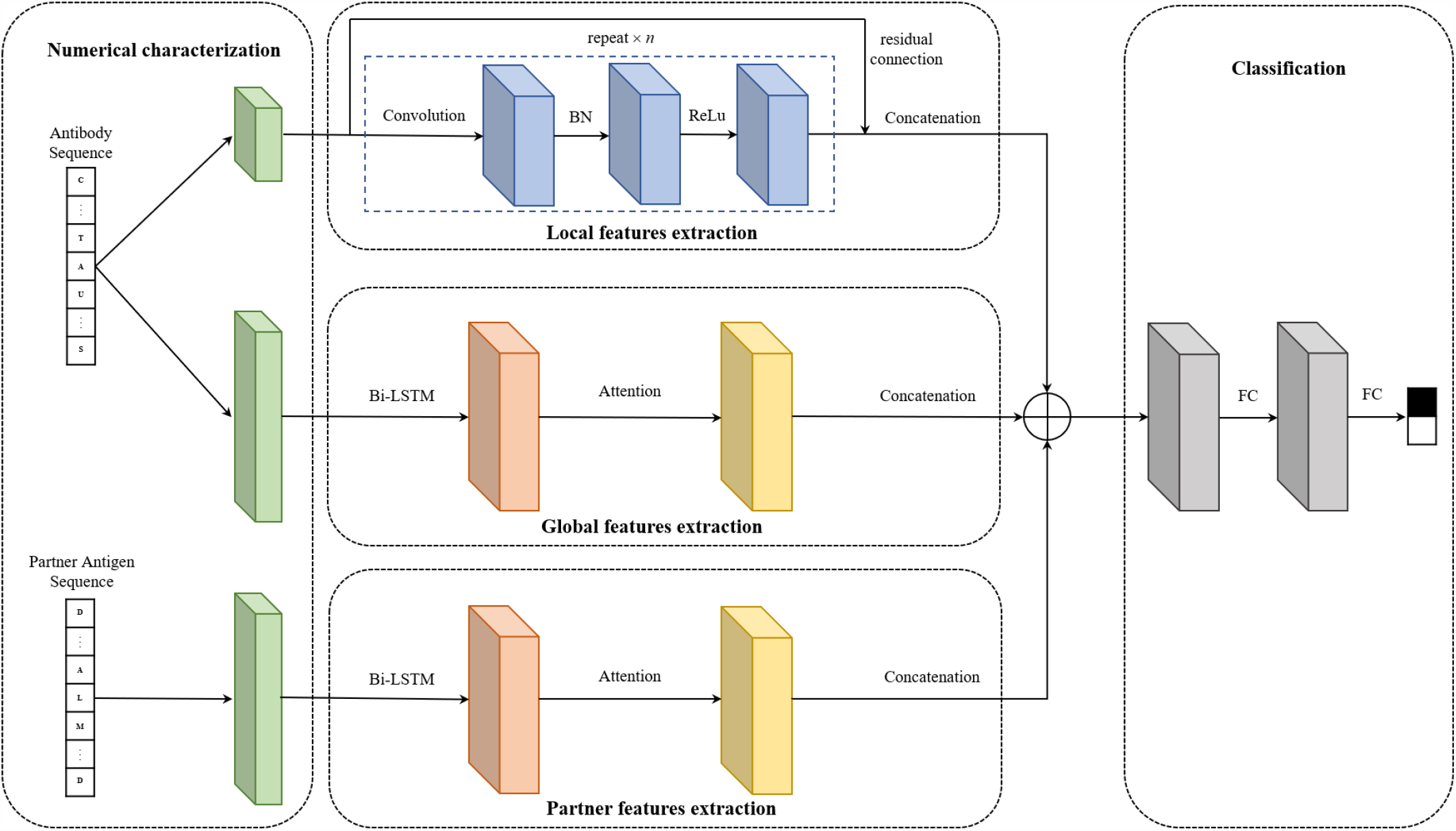
Architecture of our method

#### 2.4.1 CNNs

Convolution neural networks (CNNs) model has been used in some bioinformatic tools for protein binding site prediction[44], protein-ligand scoring[45] and protein-compound affinity prediction[46]. In our method, the input of CNNs is the local information of the target antibody residue which can be represented as *r*_*i−w*:*i*+*w*_. It means that we consider the target antibody residue at the center with 2*w* neighboring residues representing the context of target antibody residue. Those antibody residues which do not have neighboring residues in the left or right are padded by all-zero vectors. The convolutional operation is shown as:

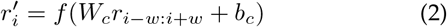

where *f* is a non-linear activation function (e.g. ReLU), *W*_*c*_ is the weight matrix, *r*_*i−w*:*i*+*w*_ the concatenation of the local information of target antibody residue, and the *b*_*c*_ is the bias vector. As Fig. 1 shows, BN means a batch norm layer and the repeat time is 3 in out model. Also, residual connection is used by adding inputs to outputs:

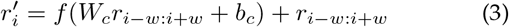

#### 2.4.2 Att-BLSTM networks

Attention-based Bidirectional Long Short-Term Memory (Att-BLSTM) networks have been used in chemical and biomedical text processing tasks[47], [48]. However, its advantage has not been exploited in biology sequence analysis such as antibody paratope prediction. In this study, Att-BLSTM networks are used to capture global features of antibody and antigen sequences.

As shown in Fig.2, the architecture of Att-BLSTM consists of four parts: input layer, Bi-LSTM layer, attention layer and output layer. The input antibody or antigen sequence is represented as a set of residues: *S* = [*r*_1_, *r*_2_, *r*_3_, ⋯, *r*_*i*_, ⋯, *r*_*l*_]^*T*^, *S* ∈ *R*^(*l*∗*d*)^

**Fig. 2.**
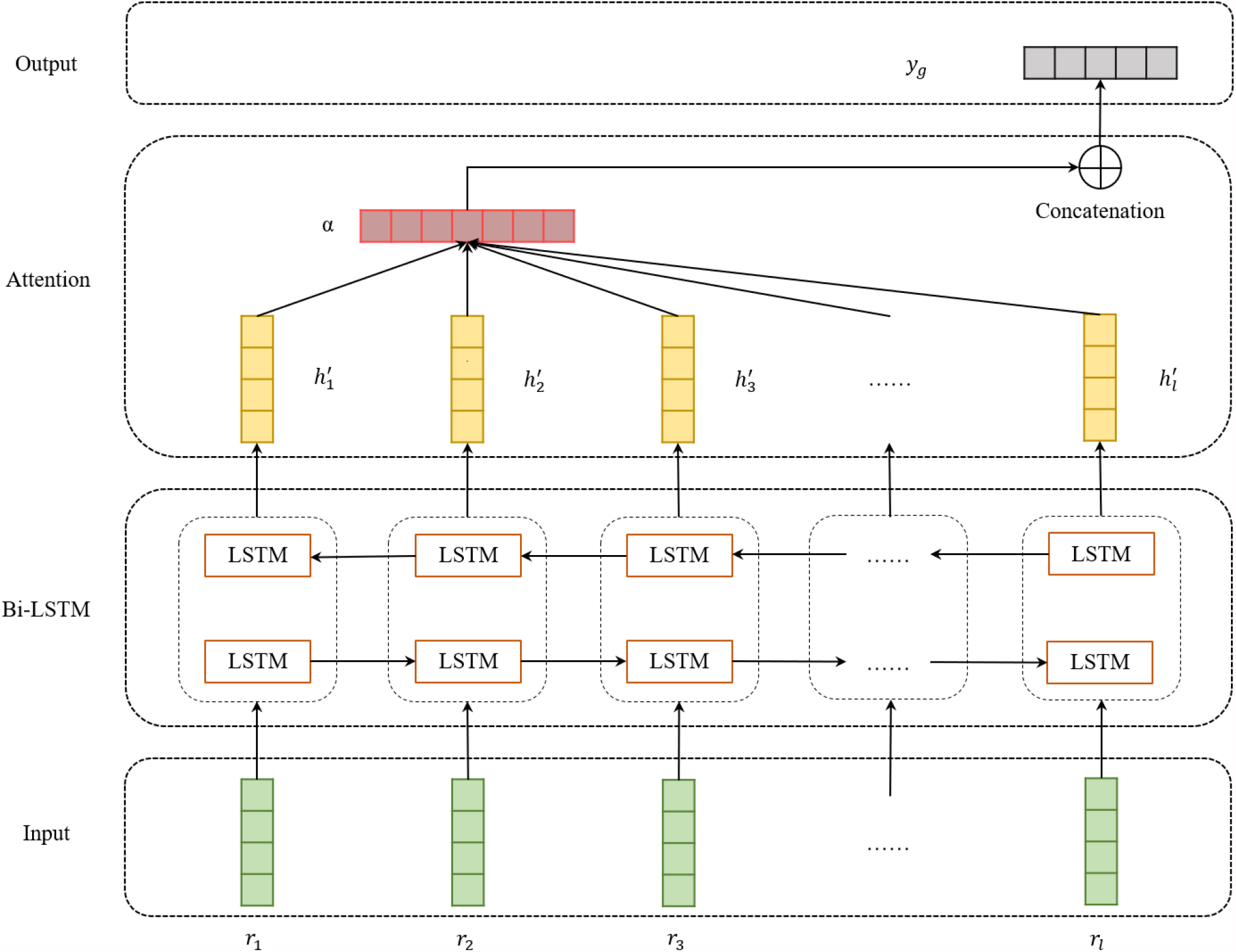
Architecture of Att-BLSTM

The Bi-LSTM layer contains two Long Short-Term Memory (LSTM) networks of which one is forward taking input residues form the beginning to the end and another is backward taking input residues from the end to the beginning. A standard LSTM contains three gates and a cell memory state to store and access information over time. Typically, a cell of LSTM can be computed at each time t as follows:

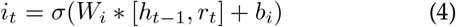

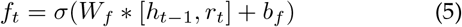

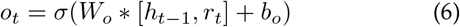

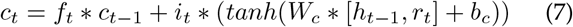

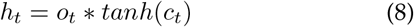

where *W*_*i*_, *W*_*f*_, *W*_*o*_ are the weight matrixs, and *b*_*i*_, *b*_*f*_, *b*_*o*_ are the biases of input gate, forget gate and output gate, respeatively. And, tanh is the element-wise hyperbolic tangent, *σ* is the logistic sigmoid function, *r*_*t*_, *h*_*t−*1_ and *c*_*t−*1_ are inputs, and *h*_*t*_ and *c*_*t*_ are outputs. For the *i − th* residue in the input antibody or antigen sequence, we combine the output 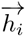 of forward LSTM and output 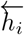 of the backward LSTM by concatenating them:

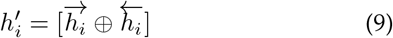

In attention layer, let *H ∈ R*^(*l∗*2*d*)^ be a input sequence 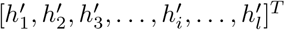 in which each element is the i-th residue output of Bi-LSTM, where l is the input sequence length, and d is the residue features dimension. The attention mechanism can be calculated as follows:

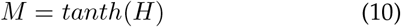

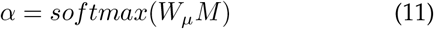

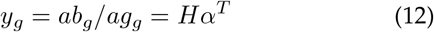

where tanh is the element-wise hyperbolic tangent, *W*_*u*_ is the weight matrix, and *α* is an attention vector. The output *y*_*g*_(*ab*_*g*_ grepresenting for the output of antibody sequence and *ag*_*g*_ representing the output of antigen sequence) is formed by a weighted sum of vectors in *H*.

#### 2.4.3 Fully-connected networks

As shown in Fig. 1 and 2, the local features extracted by CNNs is 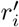 and the global features derived by Att-BLSTM networks from antibody and its partner antigen sequence are *ab*_*g*_ and *ag*_*g*_, respectively. And then, 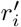, *ab*_*g*_ and *ag*_*g*_ are concatenated and fed to the fully-connected networks. The calculation of probability *y*_*i*_ for each input antibody residue belonging to paratope is shown as:

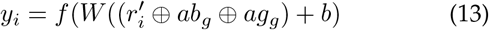

### 2.5 Training details

We implement our model using PyTorch v1.4. The training configurations are: loss function: weighted cross-entropy loss function as in [49], optimization: Momentum optimizer with Nesterov accelerated gradients; learning rate: 0.001; batch size: 32; dropout: 0.5; a fixed sliding window length: 11. The first fully connected layer has 512 nodes and the second fully connected layer has 256 nodes. Training time of each epoch varies roughly from 1 to 2 minutes depending on the global features are used or not, using a single NVIDIA RTX2080 GPU.

## 3 Results

### 3.1 Model results

To evaluate the performance of our method and other competing methods, we use the standard metrics, i.e. area under the receiver operating characteristics curve (AUC ROC), the area under the precision recall curve (AUC PR), MCC and F-score. Because our method output a probability for each antibody residue, we compute MCC and F-score by predicting residues with probability above 0.5 as paratope. To summarize the performance, all metrics are averaged over all antibodies in the testing set. We repeat the training and testing procedures five times for providing robust estimates of performance. The mean value and standard error are reported in Table 2.

**TABLE 2.** Performances comparison with competing methods

| Method | AUC ROC | AUC PR | MCC | F-score |
| --- | --- | --- | --- | --- |
| *Parapred* | 0.880 ± 0.002 | - | 0.564 ± 0.007 | 0.69 |
| *Fast-Parapred* | 0.883 ± 0.001 | - | 0.572 ± 0.004 | - |
| *AG-Fast-Parapred* | 0.899 ± 0.004 | - | 0.598 ± 0.012 | - |
| *proABC-2* | 0.91 | - | 0.56 | 0.62 |
| *Our Method* | **0.945 ± 0.001** | **0.644 ± 0.002** | **0.635 ± 0.005** | **0.819 ± 0.003** |

Table 2 shows the results of our method and other competing sequence-based antibody paratope prediction methods. Parapred uses CNNs and LSTM for predicting probability of the residues in CDRs of antibody sequence[15]. Fast-Parapred and AG-Fast-Parapred outperform Parapred by leveraging self-attention and cross-modal attention, respectively[16]. proABC-2 is based on CNNs model and takes separate heavy chain and light chain as input[14]. The results of proABC-2 is trained on the Parapred-set and uses a threshold of 0.37. As Table 2 shows, our method performs best on all metrics.

### 3.2 Effect of global features and partner features

Although global features have been proved useful in protein-protein interaction sites prediction [22] and protein phosphorylation sites prediction [23]. Those methods only use global features from self-sequence and don’t consider the partner features form the partner-sequence. To measure the effect of global features and partner features, we train our model using different feature combinations. The results are shown in Table 3. Except for AUC ROC, the combination of local-global and partner features achieves best performances on all other metrics. The dataset is imbalanced, and we consider AUC PR as the primary metric because itis more sensitive on an imbalanced dataset[50].

**TABLE 3.** Performances of our method using different features combination

| Features | AUC ROC | AUC PR | MCC | F-score |
| --- | --- | --- | --- | --- |
| Local | 0.930 ± 0.001 | 0.626 ± 0.001 | 0.608 ± 0.001 | 0.805 ± 0.001 |
| Local + Global | 0.946 ± 0.001 | 0.643 ± 0.002 | 0.633 ± 0.009 | 0.818 ± 0.005 |
| Local + Partner | **0.948 ± 0.002** | 0.632 ± 0.012 | 0.626 ± 0.018 | 0.814 ± 0.009 |
| Local + Global + Partner | 0.945 ± 0.001 | **0.644 ± 0.002** | **0.635 ± 0.005** | **0.819 ± 0.003** |

The results from Table 3 indicates that both global and partner features in our method can help for improving the model performance for antibody prediction and the combination of all features performs best. The partner antigen information is also used in AG-Fast-Parapred. AG-Fast-Parapred and Fast-Parapred are from the same work and trained on the same datasets[16].And, Fast-Parapred only utilizes antibody information. From Table 2, we can find that AG-Fast-Parapred perform better than Fast-Parapred which also show the effect of partner features.

## 4 Conclusion

In this study, we proposed a sequence-based antibody paratope prediction method leveraging local and global features of antibody and global features of partner antigen. CNNs model are used to extract the local features of antibody sequence. Att-BLSTM networks are used to capture the global features of the whole antibody and antigen sequence. We implement our model on benchmark datasets and the results show improvement of antibody paratope prediction. Moreover, our results declare that both global features from antibody and antigen are useful for performance improvement and our model performs best when both global features are used together

Though our method outperforms other competing sequence-based methods for antibody paratope prediction, it also has some limitations. First, similar like other sequence-base methods, our program takes a lot of time to generate sequence-based features by running PSIBLSAT[40] and NetsurfP-2.0[43]. Second, although the combined features improve the model performance, it is still inferior to the structure-based approaches.

In our work, we show that combing local-global and partner antigen features can be useful for antibody paratope prediction. In the future, we would focus on how to mine more structural features from antibody sequence for improving our model performance.

## Acknowledgments

This work was supported by Bingtuan Science and Technology Project(2019AB034), ‘Created Major New Drugs’ of Major National Science and Technology (No. 2019ZX09301-159), and Leading Talents Fund in Science and Technology Innovation in Henan Province(194200510002). Xiaofei Nan and Shoutao Zhang are the corresponding authors for this paper.

---

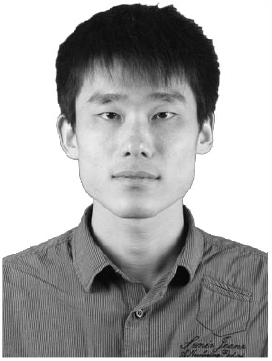

**Shuai Lu** received the BS degree from the School of Information Engineering of Zhengzhou University, Zhengzhou, Henan, China in 2014. Currently, he is working towards the PhD degree in the School of Information Engineering of Zhengzhou University. His current research interests mainly include machine learning and data mining applied to the biological field, especially in the prediction of protein-protein and antibody-antigen interactions.

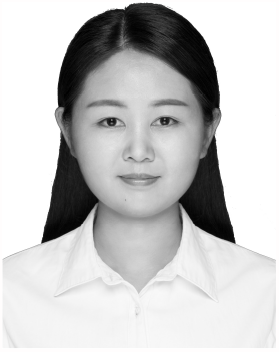

**Yuguang Li** received the MS degree from the School of Information Engineering, Zhengzhou University, Zhengzhou, Henan, China in 2018. Her research interests include bioinformatics, data mining, and NLP.

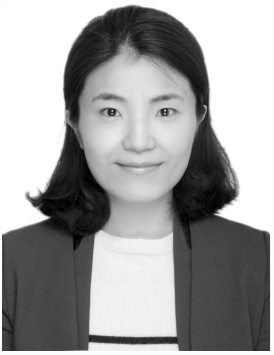

**Xiaofei Nan** received the PhD degree from the University of Mississippi. She is a vice professor in the School of Information Engineering, Zhengzhou University, Henan. Her current interests include pattern recognition, data mining, and bioinformatics.

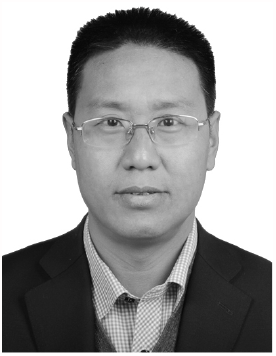

**Shoutao Zhang** received the PhD degree from the Northwest A&F University. He has published about 40 high-quality referred papers in international conferences and journals. He is a professor in the School of Life Sciences, Zhengzhou University, Henan. His research interests include bioinformatics.

